# Nuclear translocation of tagged endogenous ERK/MPK-1 MAP Kinase denotes a subset of activation events in *C. elegans* development

**DOI:** 10.1101/2021.01.22.427875

**Authors:** Neal R. Rasmussen, David J. Reiner

## Abstract

The extracellular signal-regulated kinase (ERK) MAP kinase is utilized downstream of Ras>Raf>MEK signaling to control activation of a wide array of targets. Activation of ERK is elevated in Ras-driven tumors and RASopathies, and is thus a target for pharmacological inhibition. Regulatory mechanisms of ERK activation has been studied extensively *in vitro* and in cultured cells but little in living animals. We used CRISPR to tag the 3’ end of the *C. elegans* ERK-encoding gene, *mpk-1*. Endogenous MPK-1 protein is ubiquitously expressed with elevated expression in certain tissues. We detected cytosol-to-nuclear translocation of MPK-1 in maturing oocytes and hence validated nuclear translocation as a reporter of some activation events. During developmental patterning of the six vulval precursor cells, MPK-1 is necessary and sufficient for the central cell, P6.p, to assume 1° fate. We observed MPK-1 to be recruited to the nuclei of all six VPCs in a temporal and concentration gradient centered on P6.p. This observation contrasts with previous results using the ERK-nKTR reporter of substrate activation, raising questions about mechanisms and indicators of MPK-1 activation. This system and reagent promise to provide critical insights into regulation of MPK-1 activation within a complex intercellular signaling network.

## Introduction

The mitogen-activated protein kinase (MAPK) family regulates a diverse series of cellular functions including cell proliferation, migration, and differentiation (Kolch 2005). The most well-known are the conventional MAPKs p38, JNK (c-<u>J</u>un <u>N</u>-terminal <u>k</u>inase), and ERK (<u>e</u>xtracellular signal-regulated <u>k</u>inase), all of which exhibit high degrees of conservation across metazoans (Cargnello and Roux 2011). Early work in *Drosophila* and *C. elegans* identified orthologs of ERK – rolled and MPK-1, respectively – as the terminal kinases of the Ras>Raf>MEK>ERK signaling cascade (Biggs et al. 1994; Wu and Han 1994; Lackner et al. 1994). The role of MPK-1 in *C. elegans* development was first uncovered in the vulval precursor cells (VPCs) as part of the Ras/LET-60 signaling cascade promoting 1° fate. ERK/MPK-1 was shown to be both necessary and sufficient for proper induction of 1° fate within the VPCs (Lackner and Kim 1998). ERK/MPK-1 is also a key player in other *C. elegans* tissues, including induction of excretory duct cell fate, multiple developmental events during germline proliferation, roles in nervous system function, and immune response to pathogenic bacteria (Arur et al. 2009; Church et al. 1995; Nicholas and Hodgkin 2004; Lackner and Kim 1998).

The ERK MAP kinase cascade has continued to be one of the most well studied signaling cascades due to its role as a promising pharmacological target for anti-tumor therapies in cancers with activating mutations in Ras, Raf, or upstream receptor tyrosine kinases (Ryan et al. 2015). An understanding of ERK and its mechanisms of activation has become essential, as targeted therapeutics for activated Ras and Raf have currently had limited efficacy and can promote increased activity in wild-type Raf (Durrant and Morrison 2018; Hatzivassiliou et al. 2010; Poulikakos et al. 2010).

The ERK signaling module consists of a three-tier kinase cascade with multiple phosphorylation events and negative feedback loops. Kinase activation of substrates in the linear activation cascade of Raf, MEK and ERK is largely selective, and thus can generally be considered a “signaling module”. In contrast, composition among kinases upstream of p38 and JNK is variable and exhibits significant promiscuity (Krishna and Narang 2008; Chen et al. 2011). The linearity of the Raf>MEK>ERK cascade contrasts with ERK having a large pool of substrates (∼659) and subsequent signaling outputs (Unal et al. 2017).

Upstream MAP2 kinases have the unusual property of being dual specificity kinases: they phosphorylate paired threonine and tyrosine residues adjacent to consensus docking sequences on their substrate MAP kinase (Derijard et al. 1995; Lin et al. 1995). To counterbalance activating phosphorylation of ERK, a series of <u>du</u>al <u>s</u>pecificity <u>p</u>hosphatases (DUSPs) inhibit ERK activity by dephosphorylating both the phosphorylated threonine and tyrosine residues (Huang and Tan 2012).

With ERK serving as the primary downstream signaling branch point, the field has largely relied on detection of its dual phosphorylation status to assay its activation via immunoblotting or immunostaining. This approach has revealed significant complexity in the spatial and temporal expression and activation of ERK. Nuclear translocation of ERK has also been used as indicator of its activity (Lenormand et al. 1993; Gonzalez et al. 1993), though not in model organisms. However, as active ERK and its substrates can be found both in the cytoplasm and nucleus, nuclear translocation must be interpreted with caution (Ajenjo et al. 2004; Yoon and Seger 2006; Unal et al. 2017). To allow for temporal analyses of ERK, real-time fluorescent reporters of activity like FRET (Förster resonance energy transfer) and KTR (kinase translocation reporters) have been developed for both *ex vivo* and *in vivo* contexts (Harvey et al. 2008; Regot et al. 2014; de la Cova et al. 2017). These improved tools have expanded our understanding of dynamic regulation of ERK signaling, but we still are still missing a facet of the activation of ERK: detection of endogenous ERK subcellular localization in real time.

In this study we used CRISPR/Cas9-dependent genome editing to insert sequences coding for a fluorescent::epitope tag into the endogenous *C. elegans mpk-1* gene, resulting in expressed MPK-1::mKate2^3xFlag protein. Visualization of endogenous MPK-1 – at the level of the whole animal and longitudinally throughout development – provides a novel tool to further understand its roles in signaling and development. This approach validates certain previous observations of MPK-1 expression, localization and activation in *C. elegans*. We observed tagged endogenous MPK-1 to be broadly expressed in every part of the animal and throughout development. We also detected consistently elevated expression in tissues in which MPK-1 activity has been described by other means. In agreement with previous findings, MPK-1 was expressed throughout the germline, showing brief nuclear localization in the most proximal developing oocyte. However, our approach also yielded unexpected observations. Upon induction, cytosolic-to-nuclear translocation of MPK-1 was observed in all six developing vulval precursor cells (VPCs), not just the presumptive 1° (P6.p) cell, as would have been predicted from an abundance of previous studies. Translocation in VPCs also demonstrated temporal and concentration gradients, with earliest and strongest translocation in P6.p, the VPC destined to assume a 1° fate. These unanticipated dynamics suggest that mechanisms of MPK-1 activation are more complex than previously understood. Taken together, our findings demonstrate that our tagged endogenous MPK-1 is a tool that reveals novel insights into MPK-1 regulation and roles in cell signaling complementary to those measuring phosphorylation or substrate activation.

## MATERIALS AND METHODS

### C. elegans *handling and genetics*

All strains were derived from the wild-type Bristol N2 parent strain and grown under standard conditions at 20°C unless stated otherwise (Brenner 1974). Nomenclature conforms to that of the field (Horvitz et al. 1979). All crosses were performed using standard methods, available upon request. Genotypes of strains used in this study are listed in **Supplementary Table 1**.

### Plasmids, Generation of CRISPR strains

Details of plasmid constructions are available upon request. Primers used in this study are listed in **Supplementary Table 2**, plasmids in **Supplementary Table 3**. The *mpk-1(re171[mpk-1::mkate2^SEC^3xFlag])* and *mpk-1(re172[::mKate2^3xFlag])* alleles were generated using the SEC approach of positive/negative selection for CRISPR inserts (Dickinson et al. 2015). Small guide RNAs (sgRNAs) are listed in **Supplementary Table 4**. The repair template for *mpk-1* was generated with primers oNR059, oNR060 and gBlock oNR067 for cloning into plasmid pDD285. We microinjected a mix of pCFJ104 (10 ng/µl), pNR09 (50 ng/µl), pNR10 (50 ng/µl), pNR11 (10 ng/µl) into N2 animals. Edited animals were identified by resistance to hygromycin (HygR; 5mg/ml in filtered ddH20, added directly to plates) and the Rol phenotype of the *sqt-1(*d) marker, both contained in the self-excising cassette (SEC). Homozygous animals were viable as this was C-terminal insertion. Selected animals were subsequently heat-shocked to induce expression of Cre, also contained in the SEC. Successful removal of the SEC was indicated by loss of the Rol marker (Dickinson et al. 2015). Triplex PCR detection primers oNR094, oNR095 and oNR096 were used to confirm insertion and to sequence regions of homology subject to homology-directed repair (HDR). Single and pooled animal genotyping PCR reactions used Taq PCR Master Mix (Qiagen).

### Fluorescent microscopy (imaging) and quantification of relative nuclear MPK-1 levels in VPCs

For all imaging, animals were mounted live in M9 buffer containing 2% tetramisole on slides with a 3% agar pad. DIC/Nomarski optics and fluorescence microscopy were captured using a Nikon A1si confocal microscope with 488, 561nm lasers (**Figs. 2-5; Figs. S1**,**2**) or CSU-W1 spinning-disc confocal laser microscope with 488, 561nm lasers and Photometrics Prime BSI camera (**Figs. 6, 7; Figs. S3**,**4**). Slides prepared for the spinning-disc time-lapse captured were sealed with VALAP to prevent animals from drying out. Captured images were processed using NIS Elements Advanced research, version 4.40 (Nikon). Additional deconvolution processing was performed on all time-lapse images within the Nikon Elements software utilizing the 3-D Richardson-Lucy algorithm over 35 iterations.

To determine relative levels of nuclear localization of MPK-1, animals were imaged in 10-minute intervals 24 hours post synchronization. Following deconvolution (above), fluorescent intensity measurements for both MPK-1::mKate2 and mNeonGreen::HIS-72 were recorded for each P4.p-P8.p nucleus using NIS Elements Advanced research, version 4.40 (Nikon) software. CTNF (<u>c</u>orrected total <u>n</u>uclear fluorescence) was calculated by subtracting background fluorescence. To account for variations, we then divided the MPK-1::mKate2 CTNF intensity by their corresponding mNeonGreen::HIS-72 intensity. P-values were calculated using ANOVA.

### Immunoblotting

For preparation of protein lysates, animals were washed from plates and boiled in 4% SDS loading buffer at 95°C for 2 minutes. Lysates were separated on a 4-15% SDS gel (BIO-RAD), transferred to a PVDF membrane (EMD Millipore Immobilon) and probed with the following antibodies: monoclonal mouse anti-Flag antibody (Sigma-Aldrich #F1804) and monoclonal mouse anti-α-tubulin antibody (Sigma-Aldrich #T6199) diluted 1:2000 in blocking solution overnight. Following the primary incubation, blots were incubated with the goat anti-mouse HRP-conjugated secondary antibody, (MilliporeSigma 12-349), diluted 1:5000 in blocking solution for 1hr. Imunnoblots were then developed using ECL kit (Thermo Fisher Scientific) and X-ray film (Phenix).

### Synchronized Populations

To achieve tightly synchronized populations without potential biological artifacts introduced by a bleach/starvation synchronization protocol, we utilized the NemaSync *C. elegans* synchronizer model 5000 (InVivo Biosystems). Mixed-stage animals were grown to high density on twelve to twenty 10 mm NGM plates seeded with OP50 bacteria. Animals were then washed off in M9 and added to the stabilization filter, allowing gravid adults to be separated from all other stages. Adult animals were collected from the filter and pipetted onto the harvest filter. Recently hatched L1 collected from 15-minute windows were then re-plated onto OP50-seeded plates and grown at 20°C for precisely the desired time. All reported times for time-lapse imaging refer to time post-plating of synchronized L1 animals.

## RESULTS

### Tagging endogenous MPK-1 via CRISPR

The *mpk-1* gene encodes two variants with differing promoters and hence differing 5’ ends. The longer *mpk-1b* transcript is predicted to yield a protein of approximately 51 kD, while the shorter *mpk-1a* transcript, which lacks the additional 5’ exon, is predicted to encode a protein of 43 kD (**Fig. 1A**). To ensure that we tagged both isoforms of endogenous MPK-1, we elected to insert our CRISPR tag in the 3’ end of the gene, to generate proteins tagged at the C-terminus (**Fig. 1A**). The final tagged *mpk-1* allele was confirmed through sequencing of flanking DNA and immunoblotting against the 3xFlag epitope. Both predicted isoforms were visible as expected at 77 and 69 kDa including tags, respectively (**Fig. 1B**). This reagent positions us to assay endogenous MPK-1::mKate2^3xFlag functions from a novel perspective.

**Figure 1.**
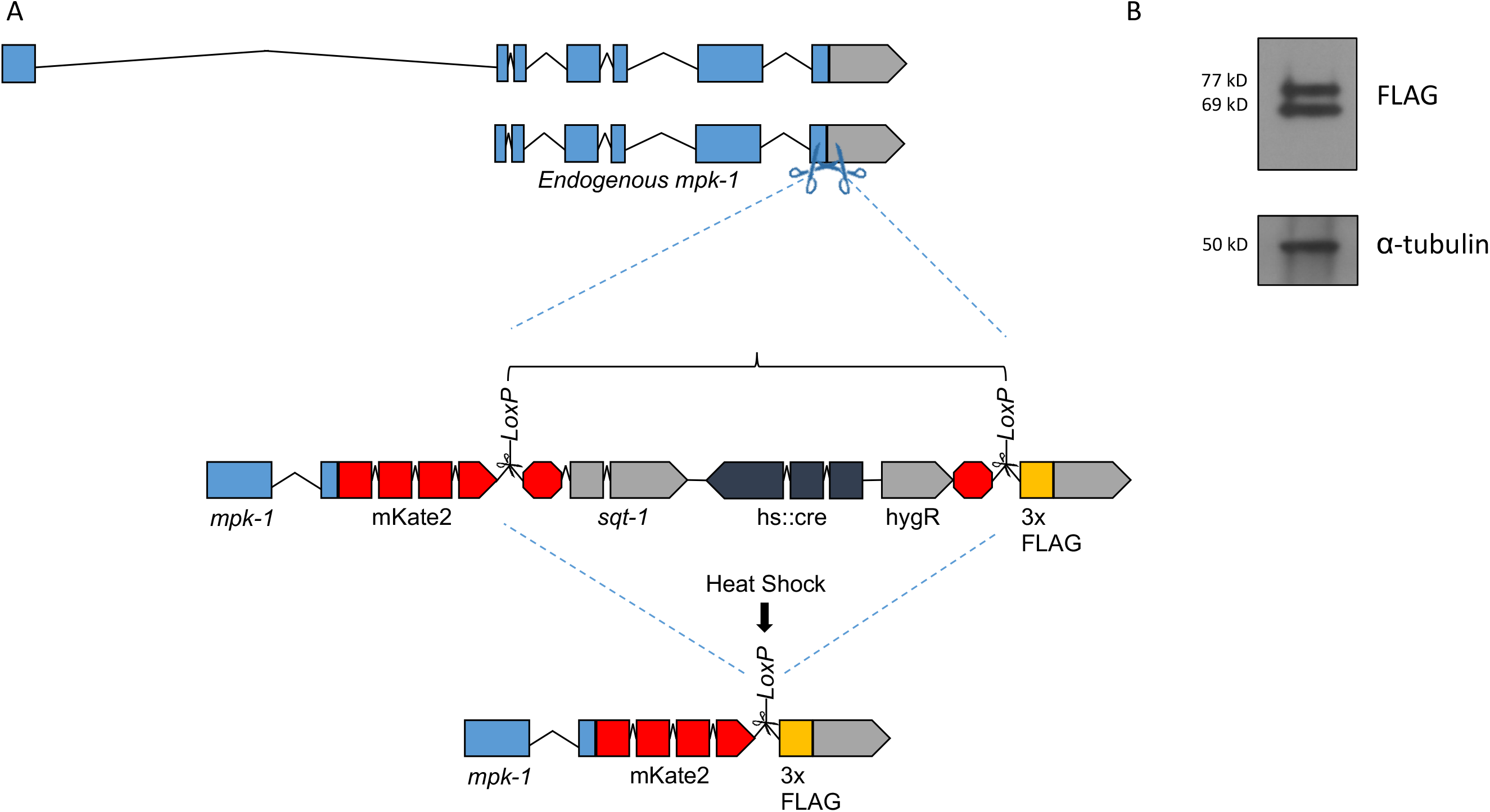
Tagging endogenous *mpk-1* using the SEC strategy. **A)** Diagram of the strategy for the CRISPR/Cas-9-dependent knock-in of mKate2::SEC^3xFlag into the 3’ end of *mpk-1*. **B)** Immunoblotting for MPK-1::mKate2^3xFlag via the 3xFlag epitope tag along with α–tubulin loading control.

### Endogenous MPK-1::mKate2 is expressed ubiquitously and throughout development

Endogenous MPK-1::mKate2 was observed to be expressed in every cell type we could discern. (**Fig. 2; S1**). We observed expression at each stage of development, from embryo to fertile adults, including the mature germline (**Figs. S1A-F; S2A-F**). Unexpectedly, expression levels were globally elevated during the L2 and L3 stages compared to L1, L4, and adults (**Figs. S1**,**S2**). It is unclear why this change might occur and be reversed. Additionally, throughout development higher levels of MPK-1 were observed in neurons in the head and in the rectal epithelium (**Fig. S1**,**S2**). We also observed elevated expression in the posterior gut, anterior gut, and pharynx (**Figs. S1**,**S2**), all consistent with pro-inflammatory functions. The highest expression levels of MPK-1 were observed in the cells in the nerve ring around the pharynx and surrounding the anus (**Fig 3A-C, G-I**).

**Figure 2.**
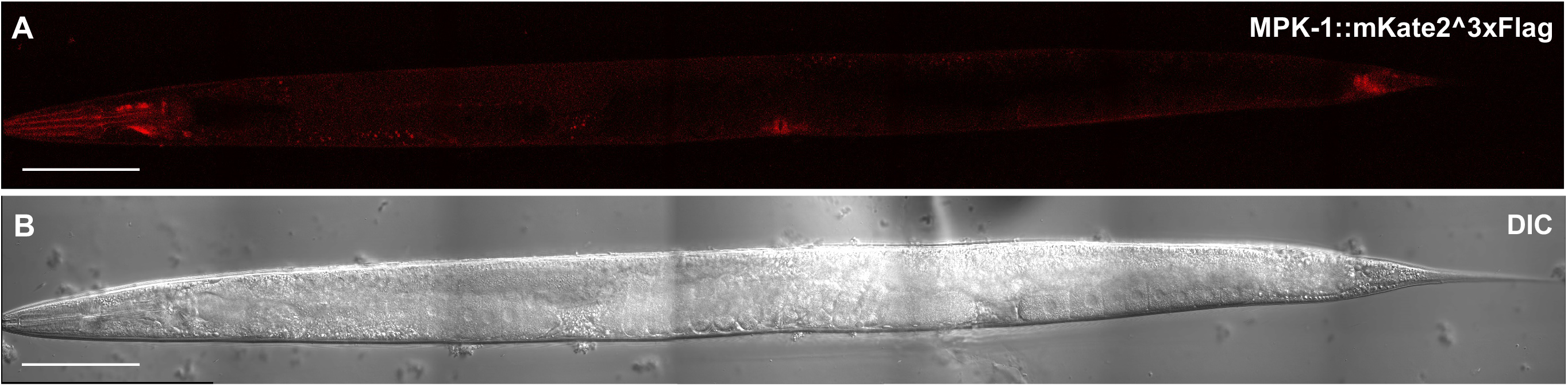
MPK-1 is expressed ubiquitously in *C. elegans*. Representative photomicrographs of **A)** MPK-1::mKate2 expression along with corresponding **B)** DIC images of an adult animal. Scale bars = 100 µm

Given the varied but broad expression of MPK-1 throughout the animal we conducted a survey of its expression in various tissues in which prior research had established a role for MPK-1. Dueto the pivotal role of MPK-1 in germline development, we first it examined its expression pattern throughout the gonad, where multiple MPK-1-dependent events are known to occur (Arur et al. 2009). Consistent with prior immunoblotting for total MPK-1 in dissected gonads, mpk-1 was expressed throughout the germline and generally localized to the cytoplasm (**Fig. 3D-F**).

**Figure 3.**
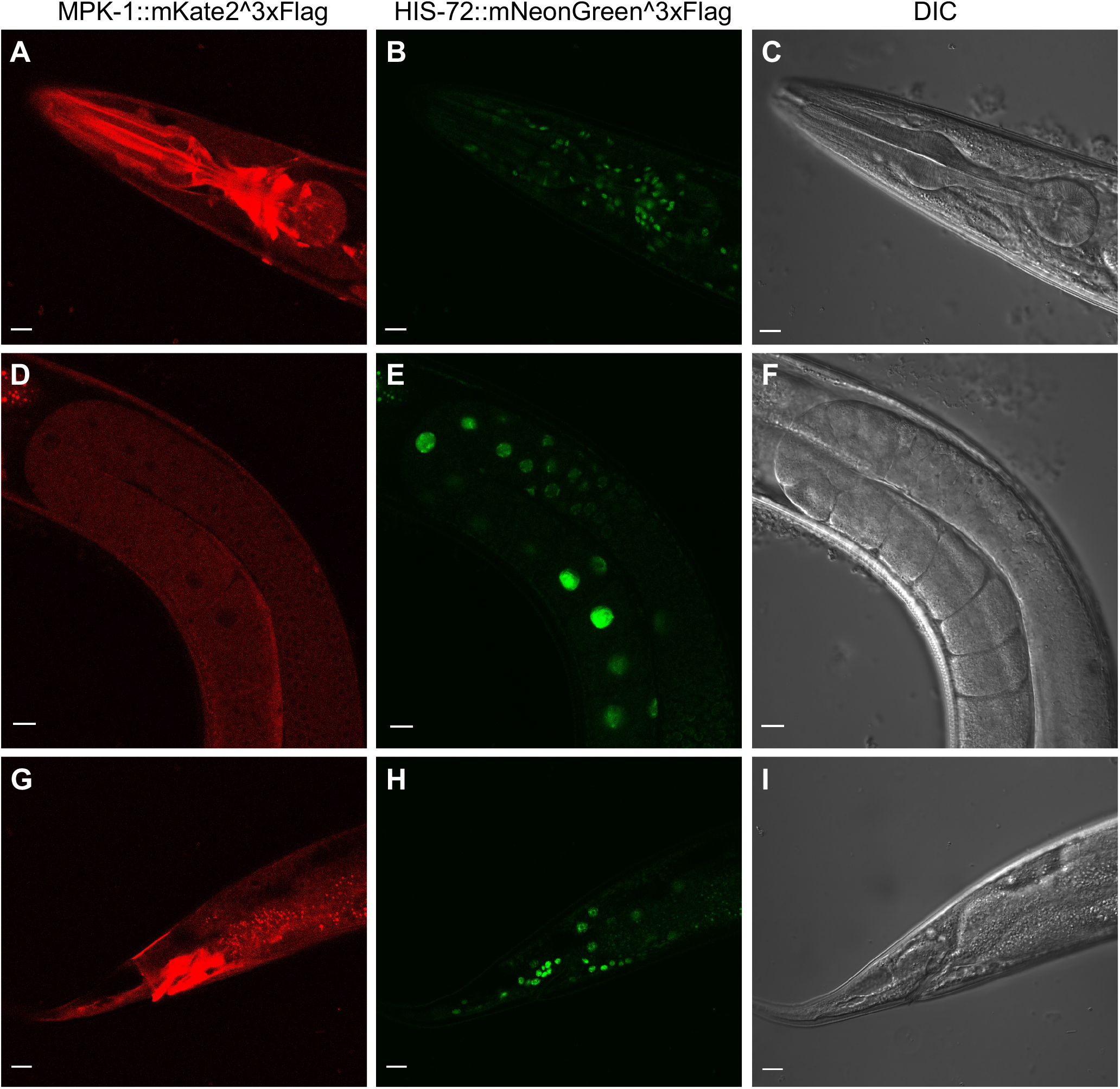
Details of tagged endogenous MPK-1::mKate2 expression and localization. Representative confocal photomicrographs are shown. **A, D, G)** Red filter set visualizing MPK-1::mKate2. **B, E, H)** Green filter set visualizing mNG::HIS-72 nuclei. **C, F, I)** DIC (Nomarksi). **A, B, C)** Adult pharynx. **D, E, F)** Mature gonad turn without embryos. **G, H, I)** Adult tail with elevated intensity in rectal epithelia. Scale bars = 10 µm.

### Endogenous MPK-1 is actively translocated to the nucleus in maturing oocytes

Substantial work has examined the role of MPK-1 signaling within the developing germline of*C. elegans* (Arur et al. 2009). This system has benefitted from its ability to be dissected from the animal for immunostaining to determine subcellular localization for both total MPK-1 and dual phosphorylated MPK-1 (dpMPK-1; Lee et al. 2007). This approach has provided snapshots to allow an understanding of the spatial and temporal expression and phosphorylation patterns of MPK-1.

We examined the expression pattern and subcellular localization of endogenous MPK-1::mKate2^3xFlag within in the proximal germline of animals 24 hrs post-mid-L4. In keeping with previous findings, MPK-1 expression was ubiquitous throughout the germline and largely excluded from the nucleus. An examination of the four oocytes most proximal to the uterus revealed infrequent nuclear translocation of MPK-1 in the most mature oocyte (position -1). Lee et al highlighted that dpMPK-1 staining and localization varied dependent upon its stage of maturation, with early oocytes displaying high nuclear expression (Lee et al. 2007). We also observation this variation in the nuclear localization of tagged endogenous MPK-1 within early oocytes (**Fig. 4**).

**Figure 4.**
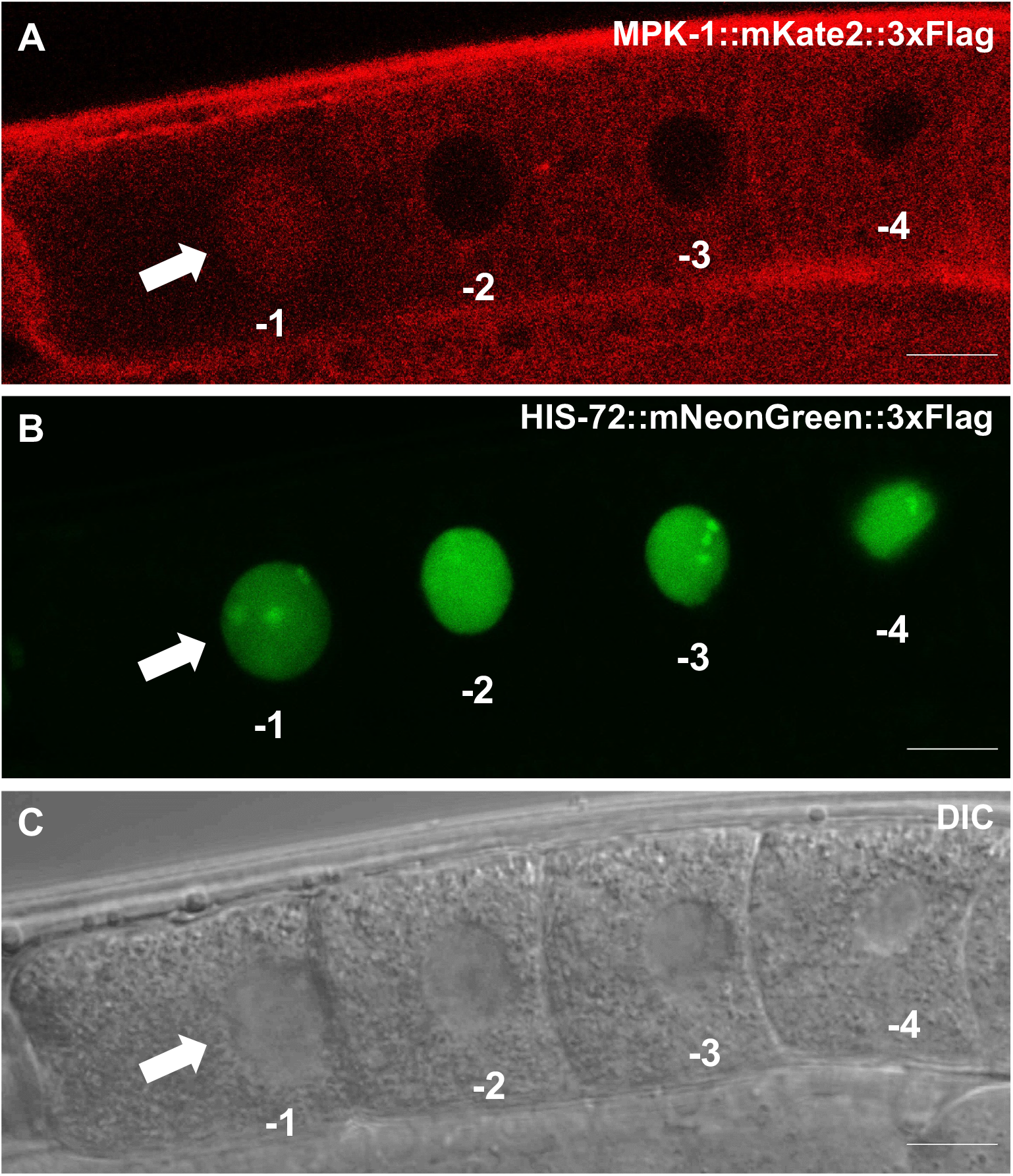
MPK-1 nuclear localization in maturing proximal oocytes. Representative confocal and DIC photomicrographs are shown. Positions are indicated as -1 (most mature) through -4 (least mature). Arrow indicates nucleus with translocated MPK-1::mK2. **A)** Red channel, MPK-1::mK2 in cytosol but with a pool of tagged protein translocated to the nucleus in the most proximal oocyte (position -1). **B)** Green channel, the same animals with green mNG::HIS-72 nuclear marker. **C)** DIC images of the four most proximal developing oocytes. Scale bars = 10 µm.

### RSKN-1 negative feedback regulates endogenous MPK-1 nuclear localization in developing oocytes

Prior work established RSKN-1, the *C. elegans* ortholog of p90 RSK kinase (Carriere et al. 2008), as a downstream target of MPK-1 that functions in a negative feedback loop to inhibit and/or restrict activation of MPK-1 in maturing oocytes: *rksn-1*-specific depletion by RNA interference resulted in expansion of the compartment of oocytes displaying dpMPK-1 staining (Arur et al. 2009). Building on our previous findings with nuclear MPK-1 expression in the most mature diakinetic oocyte, we compared animals with or without deletion of *rskn-1*. In the wild type we observed consistent nuclear exclusion of MPK-1::mKate2 in the four most proximal diakinetic oocytes (**Fig. 5A,B,E**). In contrast, in *rskn-1(ok159)* mutant animals, nuclear localization was observed in all four proximal oocytes animals (**Fig. 5C,D,F**). This result validates cytosol-to-nuclear translocation of endogenous tagged MPK-1 as a biomarker for activation.

**Figure 5.**
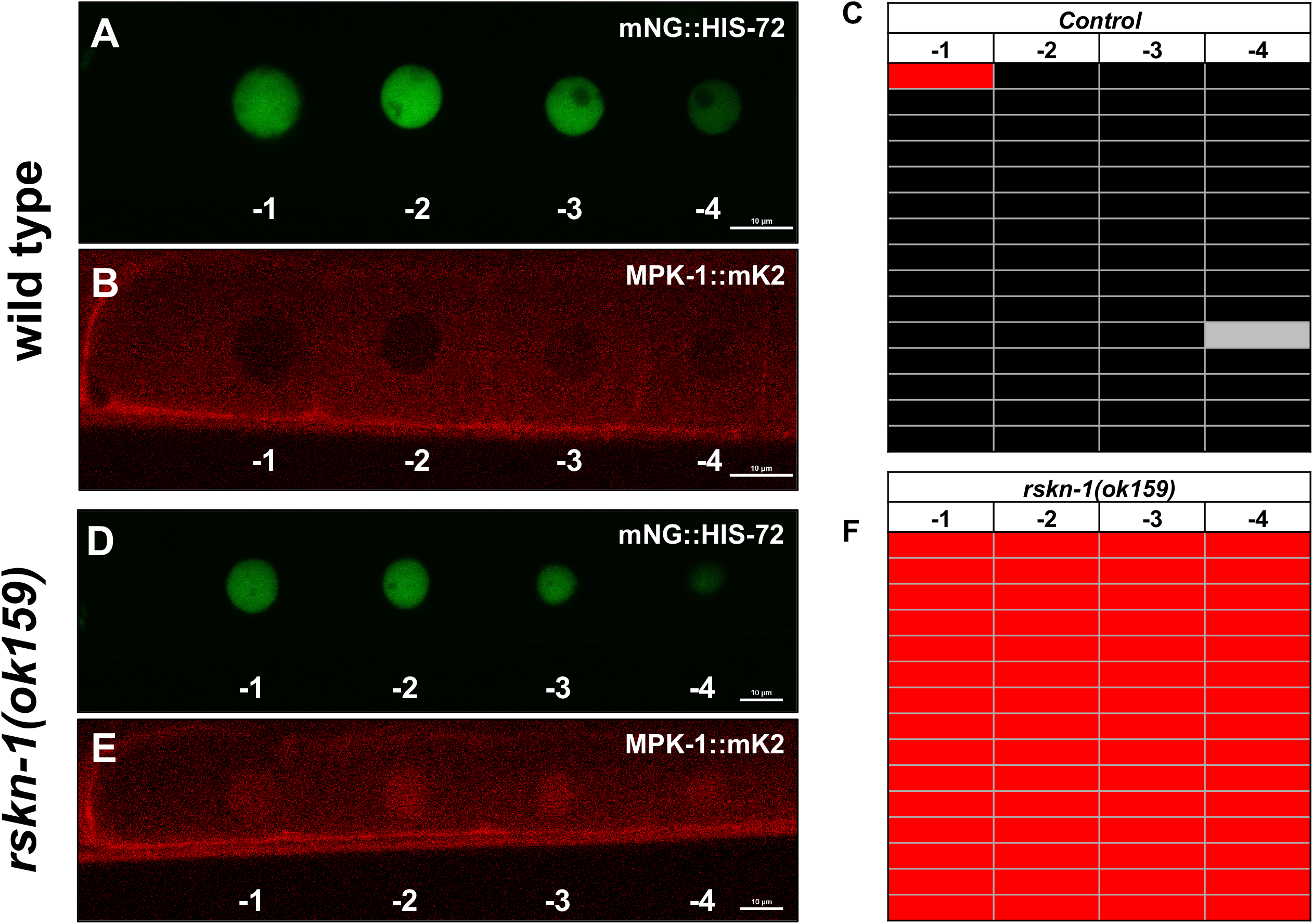
RSKN-1 negatively regulates nuclear translocation of endogenous MPK-1 in maturing oocytes. Matched representative confocal photomicrographs are shown. **A, B, C)** A wild-type animal shows infrequent translocation of endogenous tagged MPK-1 to nuclei. **D, E, F)** An *rskn-1(ok159)* deletion mutant reveals translocation of MPK-1 to every maturing nucleus. **A, D)** mNG::HIS-72-marked nuclei. **B, E)** MPK-1::mK2 signal. Scale bars = 10 µM. **C, F)** Tabulation of observed wild-type vs. mutant animals (n=15; the gray marker in **C** represents an animal in which the nucleus in position -4 was out of the plane of focus).

### Pn.p cells neighboring VPCs exhibit nuclear MPK-1 prior to VPC induction

We will describe how tagged endogenous MPK-1 translocates to the nuclei of all VPCs, the classic system for analysis of MPK-1 in *C. elegans* (see below). But first we will note our observation that P2.p, which is not a VPC and thus not competent to be induced to develop as a vulval lineage, exhibits a pool of tagged endogenous MPK-1 in its nucleus at a time when its posterior neighbors and VPCs, P3.p and P4.p, do not have nuclear MPK-1 (**Fig. 6A-C**). We propose that P2.p receives some signal through MPK-1 during a period in which VPCs are not receiving inductive signal from the AC.

**Figure 6:**
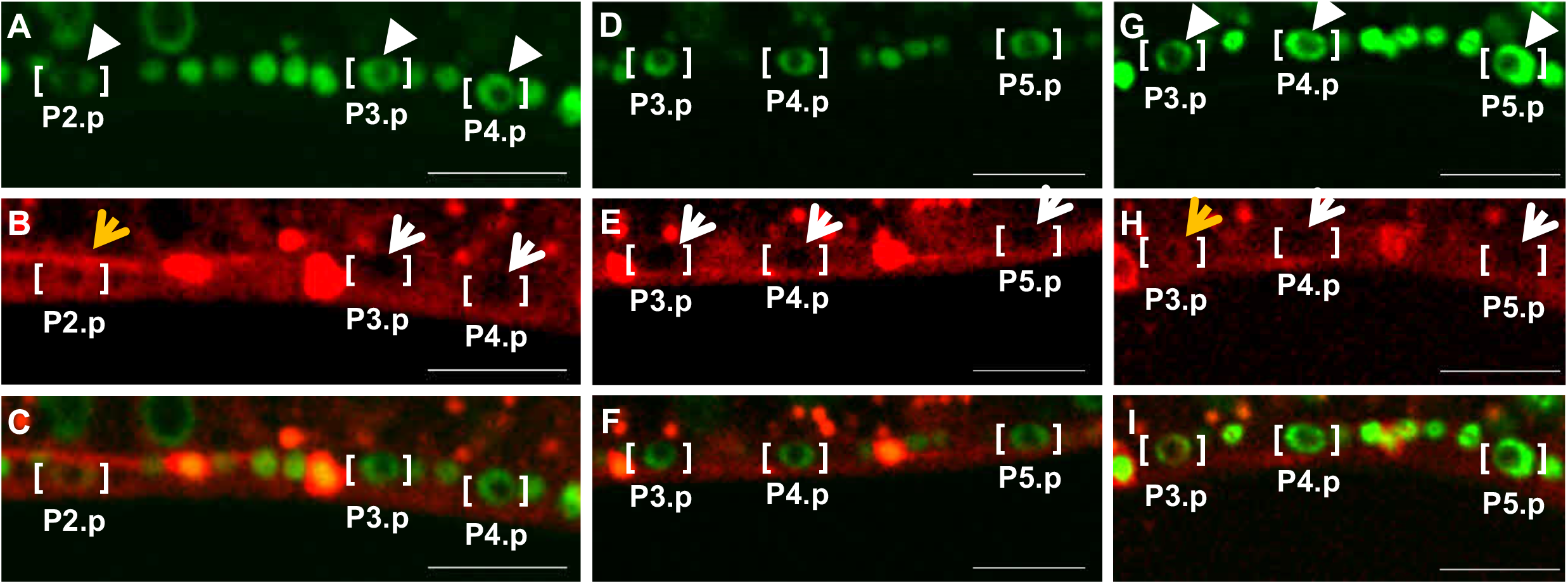
Pn.p neighbors of VPCs have high nuclear MPK-1. **A, D, G)** mNG::His-72 nuclear marker; **B, E, H)** MPK-1::mK2 red signal; **C, F, I)** Merged images. Larges spaces of signal exclusion are nuclei, small spaces of signal exclusion are nucleoli (see green DNA signal). Orange arrows indicate high nuclear MPK-1, white arrows indicate low nuclear MPK-1. The white arrowheads indicate VPC and non-VPC nuclei with differing levels of mNG::HIS-72 intensity. **A, B, C)** Nuclear MPK-1 is high in P2.p and low in P3.p and P4.p. D, E, F) Nuclear MPK-1 is low in P3.p, P4.p and P5.p. **G, H, I)** Nuclear MPK-1 is high in P3.p and low in P4.p and P5.p. Scale bars = 10 µm.

In a subset of animals, P3.p fails to achieve competence as a VPC, probably because of variable Wnt signal from the posterior during the L2, which contributes to competence, while in other animals, P3.p is a competent VPC (Euling and Ambros 1996; Myers and Greenwald 2007; Eisenmann et al. 1998). Accordingly, in some animals P3.p, P4.p and P5.p exhibit no nuclear MPK-1 (**Fig. 6D-F**), consistent with P3.p being a competent VPC in these animals and equipotent to its posterior neighbors P4.p and P5.p. In contrast, in other animals P3.p displays high nuclear MPK-1 when its posterior neighbors that are always VPCs, P4.p and P5.p, do not (**Fig. 6G-I**). We infer that these latter P3.p cells are not competent VPCs.

We observed a similar relationship between non-VPC P9.p and its competent anterior VPC neighbors P7.p and P8.p. Unlike the variability in P3.p competency, P7.p and P8.p are reported to always be competent VPCs in the wild type. We observed that at the time point where MPK-1 was excluded from the nucleus of P7.p and P8.p, a pool of MPK-1 was observed in the nucleus of P9.p (**Fig. S3**).

We do not know the nature of signals to non-VPC Pn.p cells at this stage, nor how many of the non-VPC Pn.ps respond to that signal. However, competent VPCs are presumably refractory in response to this signal, and/or non-VPC Pn.ps have a separate competency program. Additionally, absence vs. presence of nuclear MPK-1 in P3.p may be the earliest indicator of competence of P3.p as a VPC.

A phenomenon we observed was decreased expression of tagged HIS-72, an H3 histone, in non-VPC Pn.p cells. The green signal from P2.p was lower than in presumed competent P3.p and P4.p (**Fig. 6A**), including in P3.p where elevated nuclear translocation of tagged endogenous MPK-1 suggested that the cell was non-competent as a VPC (**Fig. 6G**). Similarly, the nuclear mNG::HIS-72 signal of non-VPC P9.p was weaker than its lineal homologs P7.p and P8.p (**Fig. S3A**). We speculate that HIS-72 is expressed at higher levels in VPCs than in non-VPC Pn.p cells to confer differential regulation of gene expression.

### Cytosol-to-nuclear translocation of endogenous MPK-1 is observed in all VPCs

We examined cytosolic-to-nuclear translocation of tagged endogenous MPK-1 in developmental patterning of vulval precursor cell (VPC) fates. MPK-1 was originally identified in the VPC system by the ability of reduction-of-function mutations in *mpk-1* to suppress the ectopic Multivulva (Muv) phenotype conferred by constitutively activated LET-60/Ras (Wu and Han 1994; Lackner et al. 1994). Subsequent studies implicated MPK-1 in phosphorylation and repression of the downstream transcription factors LIN-1/ETS and LIN-31/FoxB, which coordinate expression of the LIN-39 Hox transcription factor and the Mediator Complex to induce 1° fate (Jacobs et al. 1998; Fantz et al. 2001; Wagmaister et al. 2006b; Wagmaister et al. 2006a; Bagshaw 1993; Tan et al. 1998; Miller et al. 1996; Underwood et al. 2017).

We failed to observe nuclear translocation of endogenous MPK-1 using the point-scanning Nikon A1 confocal microscope using randomly selected L3 stage animals. We reasoned that a combination of signal too faint to detect and too transient to encounter with regularity among randomly selected L3 animals compromised our efforts at detection. Consequently, we turned to a spinning disk confocal microscope with a more sensitive detector. We also used L3 animals synchronized by the NemaSynch filtration device, without the use of hypochlorite treatment and starvation that is typical in the field (see Materials and Methods). With these approaches, we were able to observe exclusion of MPK-1 from VPC nuclei followed by translocation of protein into VPC nuclei. However, detection of nuclear translocation of MPK-1::mK2 was still at the limit of detection of the instrument, and required deconvolution software to visualize (see Materials and Methods).

Since MPK-1 is necessary and sufficient for induction of 1° fate, we reasonably expected to observe nuclear translocation of tagged endogenous MPK-1 only in P6.p, the presumptive 1° cell. Unexpectedly, we observed translocation of MPK-1 to the nucleus in all six VPCs (**Fig. 7**). Our observation indicates that assessment of MPK-1 activation by nuclear translocation is qualitatively different than assessment by the previously reporter ERK-nKTR reporter (de la Cova et al. 2020).

**Figure 7:**
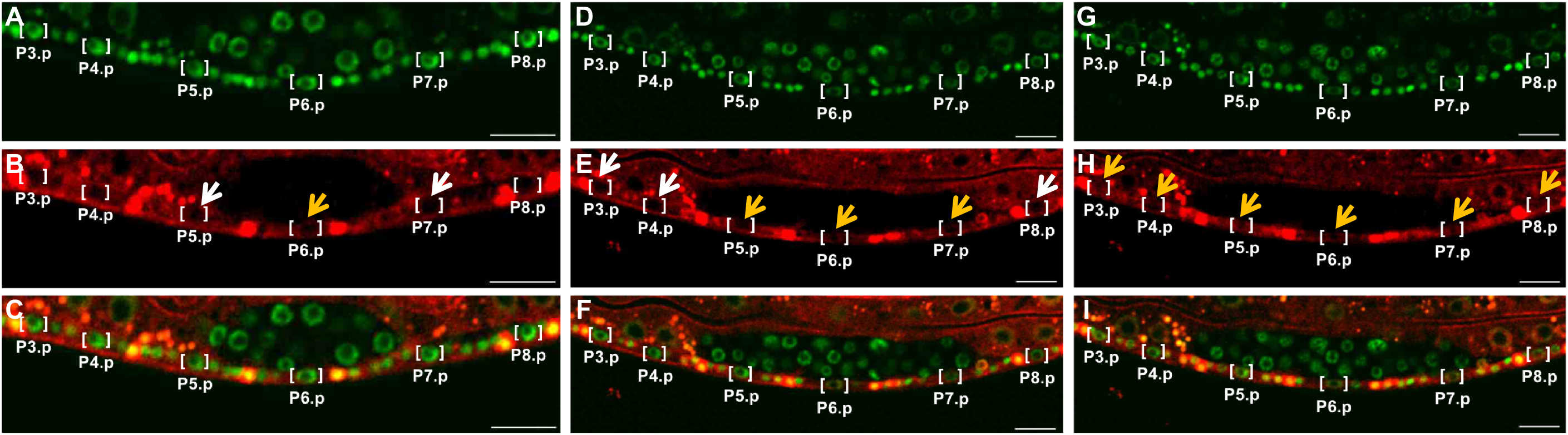
MPK-1 enters the nuclei of all VPCs, starting with P6.p. Confocal photomicrographs of tagged endogenous MPK-1 and HIS-72. **A, D, G)** Green channel mNG::HIS-72. **B, E, H)** Red channel MPK-1::mKate2. **C, F, I)** Merged images. Orange arrows indicate nuclear MPK-1 signal, white arrows indicate nuclear exclusion of signal, judged primarily by visualization of the nucleolus in the red channel. Left-to-right are three time points. We were unable to reproducibly image one animal continually throughout the time course of VPC induction, due to indicators of desiccation and toxicity. In this series of confocal photomicrographs, the first set of panels (**A-C**) is a different animal than in the second and third sets (**D-F** and **G-I**), which are the same animal at different time points. But these figures are representative of the observed process. The animal in **A-C** is also enlarged relative to later stages to better see VPC nuclei. Scale bars = 10 µm.

### Translocation in VPCs occurs earlier and at higher level in P6.p than in flanking VPCs

The cytosol-to-nuclear translocation of MPK-1::mK2 occurred in a temporal gradient centered on the source of signal, the anchor cell: first in the Pn.p cell closest to the AC, P6.p (the presumptive 1° cell; **Fig. 7A-C; Fig. S4**), then in P5.p and P7.p (presumptive 2° cells **Fig. 7D-F**), and lastly in P3.p, P4.p and P8.p (presumptive 3° cells; **Fig. 7G-I**; **Supplementary Movie 1**). This observation indicates that all six VPCs receive the inductive signal from EGF/LIN-3.

MPK-1::mK2 is also recruited to P6.p, the presumptive 1°, at higher levels than surrounding presumptive 2° and 3° cells. After nuclear translocation was completed in all VPCs, we graphed levels of nuclear MPK-1::mK2 as a ratio to mNG::HIS-72. P6.p harbors significantly higher nuclear MPK-1::mK2 than do the neighboring P5.p and P7.p (P = 0.02 and 0.05 respectively) (**Fig. 8A**), or relative to P4.p and P5.p (P = 0.05; **Fig. 8B**). Consequently, we conclude that MPK-1::mK2 translocation into VPCs occurs in both a temporal and a concentration gradient centered on P6.p, the presumptive 1° cell. This temporal gradient of nuclear entry of MPK-1::mK2 into VPCs appears to mirror the hypothetical morphogen gradient of EGF inferred from genetic analyses (Sternberg and Horvitz 1986; Katz et al. 1995; Katz et al. 1996).

**Figure 8:**
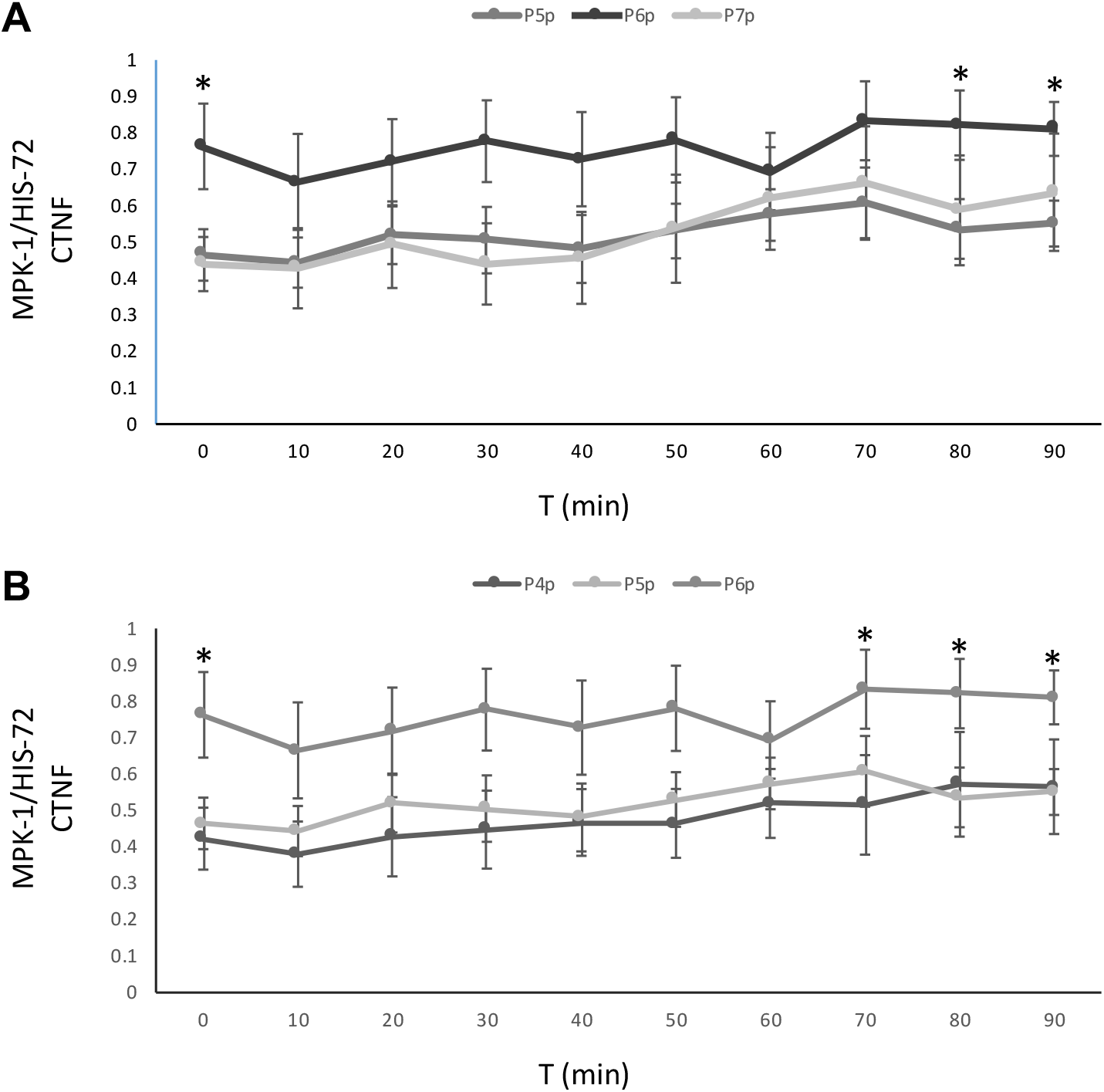
MPK-1 Nuclear Localization is higher in P6.p. **A)** Quantification of nuclear MPK-1::mK2, determined by the ratio of the CTNF (<u>c</u>orrected total <u>n</u>uclear fluorescence) between MPK-1::mK2 and mNG::HIS-72 is shown for P5.p, P6.p and P7.p. (Standard error was calculated for each time point across the samples, n=6). **B)** Quantification of nuclear MPK-1::mK2, determined by the ratio of the CTNF between MPK-1::mK2 and mNG::HIS-72 is shown for P4.p, P5.p and P6.p. (Standard error was calculated for each time point across the samples, n=6). * = P-value <0.05 as calculated by ANOVA

Taken together, our observations of the behavior of endogenous tagged MPK-1 in VPCs suggest unexpectedly rich information encoded in the Ras>Raf>MEK>ERK signal in time and space. This information may reflect the LIN-3/EGF morphogen gradient from the Anchor Cell, lateral signaling via LIN-12/Notch as part of sequential induction of VPCs, or as-yet unknown feedback loops or parallel signals that regulate the MPK-1 signal at the level of cytosol-to-nuclear translocation.

## DISCUSSION

By tagging the endogenous *C. elegans* ERK/MPK-1 and tracking its expression and subcellular movements *in vivo*, we have obtained a unique perspective on activation of MPK-1 by upstream signals. We emphasize that this is only one view of ERK activation: antibody detection of the dpMPK-1 phosphorylation or use of ERK-nKTR reporter of substrate phosphorylation are yet other, complementary views, though the former may not be feasible in the VPCs. But taken together these reagents can lead us to a thorough understanding of ERK activation and provide tools to interrogate mechanisms governing this phenomenon.

As anticipated, we observed that tagged endogenous MPK-1 is expressed ubiquitously. Expression is elevated in the nerve ring, perhaps a reflection of density of axonal/dendritic projections in ganglia of neurons (Chen et al. 2011). And we observed elevated expression in the rectal epithelium, a site of action that has been associated with anti-inflammatory activity in response to a nematode-specific bacterial pathogen, *Microbacterium nematophilum*, but presumably also in preparation for defense against other pathogens (Hodgkin et al. 2013; Anderson et al. 2013). Elevated expression via extrachromosomal array was previously observed in the rectal epithelium under control of the EGL-5 homeobox transcription factor, providing developmental insight into regulation of MPK-1 expression in different tissues (Nicholas and Hodgkin 2009), while here we observe the same phenomenon with endogenous protein. Expression in the anterior gut, posterior gut and pharynx may also be indicators of an inflammatory mechanism to protect against pathogens.

Endogenous MPK-1 is also expressed throughout the germline. Tagged endogenous MPK-1 allowed us to validate nuclear translocation of MPK-1 as a readout of upstream activation. Occasional proximal-most oocytes reveal nuclear MPK-1, and this field is expanded in animals mutant for *rskn-1*, a downstream kinase of MPK-1 possibly serving as a negative feedback loop (Arur et al. 2009). In the VPCs, in which dpMPK-1 has not been evaluated due to difficulty of fixation of somatic vs. germline structures, we observed translocation of MPK-1 into all six VPC nuclei.

### MPK-1 cytosolic-to-nuclear translocation in the VPCs

The VPCs are a complex system in which at least four signaling cascades are orchestrated to generate the 3°-3°-2°-1°-2°-3° pattern of cell fates with fidelity: core 1°-promoting Ras>Raf>MEK>ERK (Sundaram 2013) and 2°-promoting Notch (Chen and Greenwald 2004) signals, coupled with modulatory 1°-promoting PI3K>PDK>Akt (Nakdimon et al. 2012; Shin et al. 2019) and 2°-promoting Ras>RalGEF>Ral>Exo84>GCK-2>MLK-1>PMK-1 signals (Shin et al. 2018; Zand et al. 2011). Additionally, temporal control of VPC patterning is tightly coordinated at the level of the entire animal, likely with the heterochronic system and cell cycle (Wang and Sternberg 1999; Ambros 1999; de la Cova et al. 2020). We observed cytosolic-to-nuclear translocation of MPK-1 in all six VPCs in the window during which the VPCs are patterned by LIN-3/EGF. We also observed both spatiotemporal and concentration gradients of MPK-1 translocation to the nucleus, focused on P6.p. Our observations are reminiscent of the graded morphogen signal inferred from classic developmental experiments in the vulva (Sternberg and Horvitz 1986; Katz et al. 1995; Katz et al. 1996). Are the temporal and concentration gradients of MPK-1 translocation, centered on P6.p, a direct reflection of a gradient of growth factor activation of its receptor, LET-23/EGFR? Alternatively, the gradients we observed may reflect the interplay of signals active in naïve VPCs, or in parallel to the Ras>Raf>MEK>ERK signal. Our results suggest that regulation of MPK-1 in VPCs is complex, and likely to be subject to a gating phenomenon that restricts the activity to P6.p, at precisely the correct time to induce 1° fate.

### Regulation of MPK-1/ERK activation

Other signaling axes would naively be expected to extinguish sustained activation of MPK-1 in cells other than P6.p. For example, the lateral signal from LIN-12/Notch, in which MPK-1-dependent synthesis of redundant DSL ligands activate LIN-12/Notch in neighboring VPCs P5/7.p to assume 2° fate (Chen and Greenwald 2004), might be predicted to preclude MPK-1 activation in those cells, or at least activation of substrate transcription factors. A transcriptional client gene of LIN-12/Notch, the ERK phosphatase LIP-1, is expressed in P5/7.p as a consequence of lateral signal (Berset et al. 2001). Signaling from LET-23/EGFR is similarly thought to be repressed by a receptor tyrosine phosphatase, DEP-1, thus further restricting ERK activation to P6.p as a consequence of initial sequential induction (Berset et al. 2005).

In contrast, we observe that endogenous MPK-1 enters the nuclei of all six VPCs, which implies that all six receive inductive signal via the LET-60/Ras>LIN-45/Raf>MEK-2/MEK MAP kinase cascade. In contrast to our results with tagged endogenous MPK-1, the ERK-nKTR reporter of ERK activation of substrates suggested that MPK-1 was activated only in P6.p during the L3 stage. (The ERK-nKTR marker was also active in pulsatile waves prior to induction, in the L2 stage, hinting that some form of “pre-patterning” occurs prior to induction occurs; de la Cova et al. 2017). Though ectopically expressed, this reporter is single copy and hence unlikely to be subject to undesirable effects of over-expression, and is well validated elsewhere in the animal. Furthermore, activation of the ERK-nKTR biomarker closely resembles what we would expect from such a reporter predicted by the genetics: activation restricted to P6.p during the window in which P6.p is induced by the Ras>Raf>MEK>ERK cascade.

In addition to contradiction by the results using the ERK-nKTR reporter, our observation is conflict with transcriptional reporter analysis of genes reported to be responsive to MPK-1 1°-promoting signaling. During VPC induction, the promoter of the *egl-17* gene drives expression of CFP in an abbreviated gradient: expression is strong and sustained in P6.p but weak and transient in P5/7.p. The transient expression in P5/7.p is a result of LIN-12/Notch-dependent expression of the LIP-1/ERK phosphatase and other lateral signaling target genes (Yoo et al. 2004). Another reporter, the promoter of *lag-2* driving expression of YFP, is expressed only in 1° lineages (Zhang and Greenwald 2011).

We surmise that the discrepancy between the two methods of measurement reveals overlapping systems of regulation of MPK-1 to keep its activation constrained both spatially and temporally. At least at the level of MPK-1 translocation, activation of MPK-1 appears to be independent of LIN-12/Notch or expression of LIP-1. So there must be at least one other mechanism for gating the activity of MPK-1. Could output of MPK-1 be regulated through a series of interactions with transcription factors? This interpretation is unlikely, given that ERK-nKTR is a reporter of direct phosphorylation of substrate, and thus reflects activation of MPK-1 upstream of transcription factors (de la Cova et al. 2017; Regot et al. 2014). But possibly the ERK-nKTR reporter does not represent dpMPK-1, just the ability to phosphorylate a specific, defined substrate. Or perhaps MPK-1 is rapidly dephosphorylated in all but P6.p though is not accompanied by nuclear export, but signaling to downstream LIN-1 and LIN31 transcription factors is nonetheless incapacitated.

### General MAP Kinases

Other subfamilies of MAP kinases may share regulatory mechanisms. Our lab documented that tagged endogenous p38/MAP kinase, the known endpoint of the Ral 2°-promoting modulatory signal, is partially nuclear in every somatic cell of the animal (Shin et al. 2018; expression was likely silenced in the germline). This result was not observed with transgenic over-expression of GFP-tagged PMK-1 (Mertenskotter et al. 2013; our unpublished results), perhaps due to unfavorable signal-to-noise ratio attendant upon over-expression. Like ERK, p38 MAP kinases are also expected to undergo nuclear translocation upon activation (Ben-Levy et al. 1998). But perhaps a live animal experiences tonic, low-level activation of inflammatory responses mediated by PMK-1, maybe as a preventative measure against an environment with many pathogens. Thus, a small pool is always recruited to nuclei, but this pool may be miniscule compared to over-expressed heterologous protein, and so is lost due to infelicitous signal-to-noise ratio.

Still other subfamilies of MAP Kinases use nuclear translocation as a step in their activation of nuclear targets. JNK MAP Kinases translocate upon activation (Liu et al. 1996). The fourth subfamily of MAP Kinases, the non-canonical ERK5 with its transactivation domain that regulates transcription, is also regulated by translocation to the nucleus upon activation by upstream cascades (Gomez et al. 2016). Thus, this regulatory modality of MAP kinases is well-established.

Importantly, we note that many MPK-1 targets are not transcription factors or other proteins occupying the nucleus, and thus are not subject to nuclear translocation of MPK-1 as a biomarker for activation. This is clearly true for MPK-1 substrates in the germline, where myriad non-nuclear substrates have been identified and dpMPK-1 detected via antibody staining of fixed, extruded gonads (Arur et al. 2009). But non-nuclear substrates throughout the animal are also likely to be subject to phosphorylation by MPK-1. This phenomenon restricts the utility of tagged endogenous MPK-1 to events in the nucleus.

Our analysis points to nuclear translocation of endogenous MPK-1 as a robust system for analyzing activation in a live animal. This includes transient activation during developmental patterning of VPC fate, which was detected only at the lower limit of sensitivity of our instrument. We also conclude that the VPCs are a system where multiple levels of regulation of ERK are employed to achieve the desired developmental outcome. Though outside the scope of this analysis, nuclear translocation of endogenous MPK-1, the ERK-nKTR reporter, and perhaps antibody detection of dpMPK-1 could be deployed in concert to disentangle relationships between different modalities of regulation of ERK activation in an active developmental system. In the end there can be only one: P6.p assumes 1° fate in 99.8% of wild-type animals (Braendle and Felix 2008; Shin et al. 2019). Layers of regulation of ERK, including control of nuclear translocation, may impose strictures that contribute to this level of developmental fidelity.

## ACKNOWLEDGEMENTS

This work was supported by NIH grant R01GM121625 and John Templeton Foundation Grant ID# 61099 to D.J.R., and ACS PF-16-083-01 post-doctoral fellowship to N.R. Some strains were provided by the CGC, which is funded by NIH Office of Research Infrastructure Programs (P40 OD010440). Wormbase was used regularly. We thank Nikon representatives for providing expertise, resources, and the instrument demo in which we first observed the phenomenon of MPK-1 translocation in VPC nuclei.

---

**Supplemental Movie 1:** Time lapse film of MPK-1::mK2 entering P6.p first. Arrows indicate P6.p and other VPCs.

**Supplementary Table 1.**
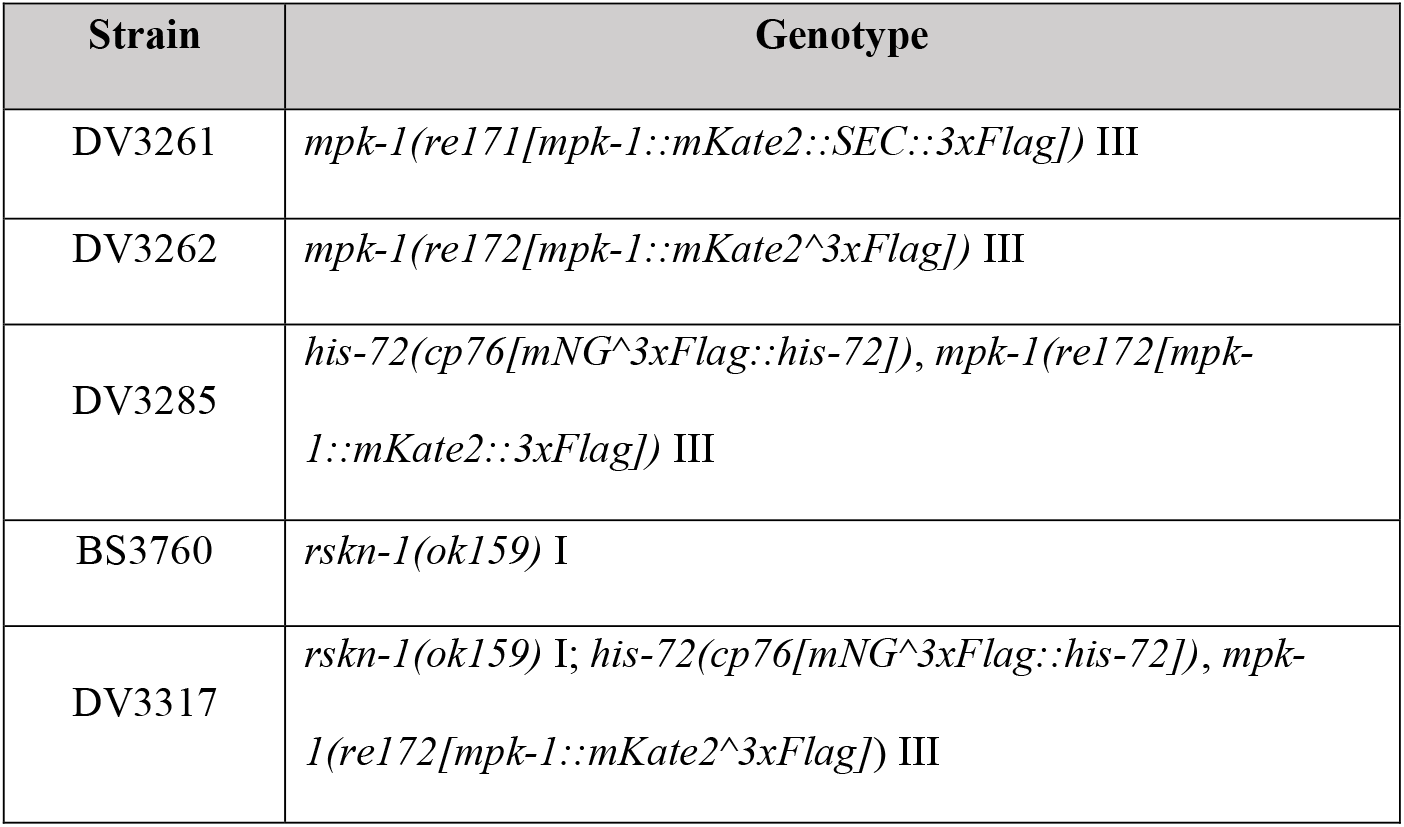
Strains.

**Supplementary Table 2.**
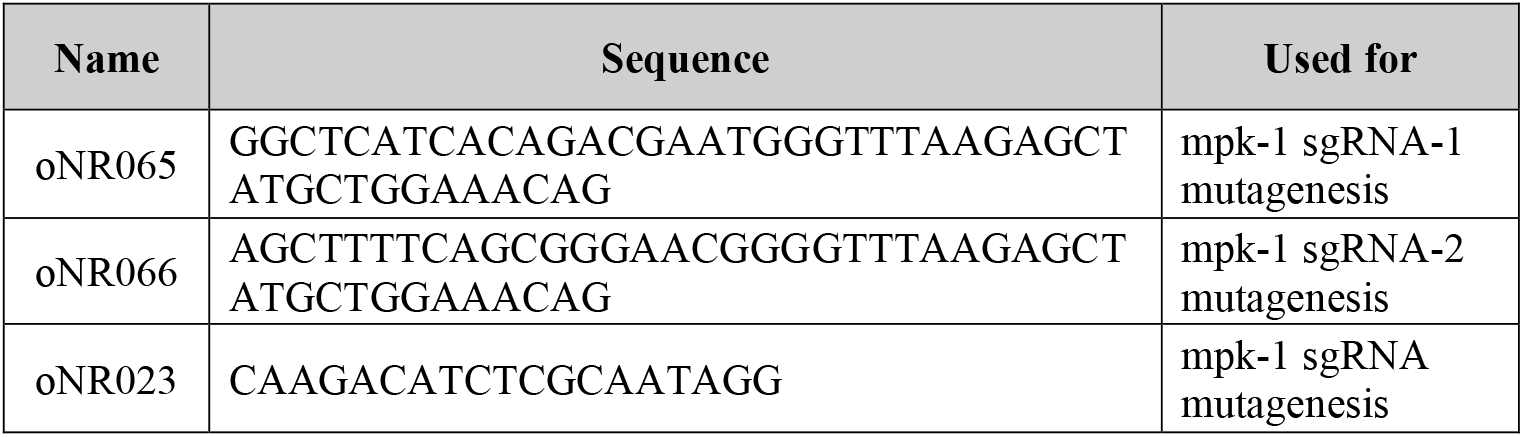
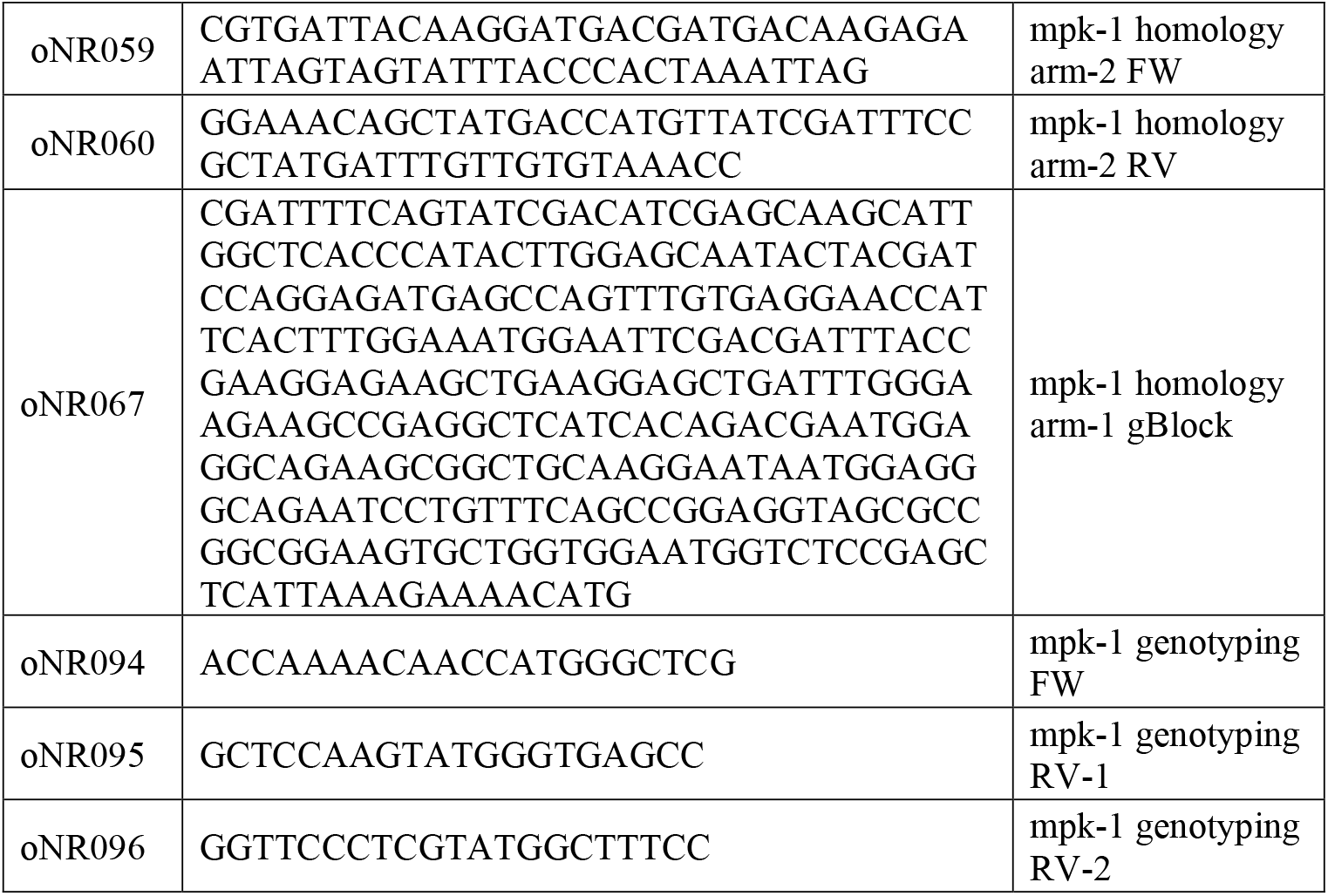
Primers.

**Supplementary Table 3.**
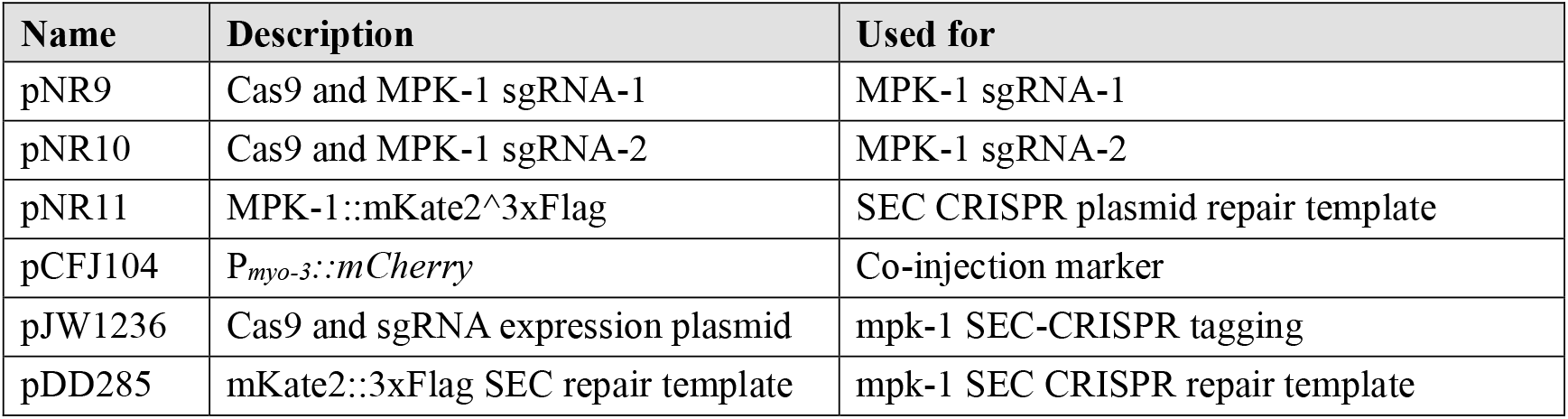
Plasmids.

**Supplementary Table 4.**
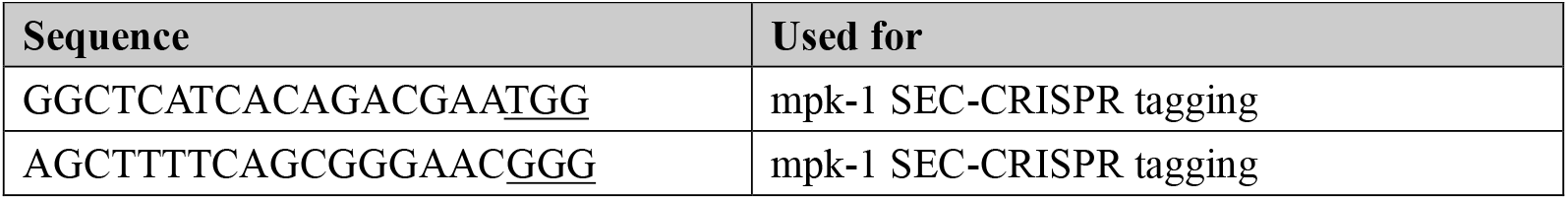
sgRNA sequences and PAMs.

